# The network structure of cross-feeding impacts microbial community diversity under growth-inhibiting stresses

**DOI:** 10.1101/2024.10.29.620519

**Authors:** Daniel P. Newton, Po-Yi Ho, Kerwyn Casey Huang

## Abstract

Perturbations such as dietary shifts and drug treatment can reduce gut microbiome diversity, with negative health effects on the host, but predicting changes to community diversity is challenging due to the many mechanisms by which species interact. While both nutrient competition and cross-feeding can play major roles in microbiota assembly, the effect of environmental stresses such as antibiotics is typically studied experimentally in monoculture, and most theoretical studies have focused on communities that interact via nutrient competition alone. To investigate the interplay between nutrient competition, stresses that inhibit growth, and cross-feeding, we introduce a consumer-resource model that includes all three factors and captures a wide range of cross-feeding network architectures with a single parameter. For three-species communities, we found that coexistence during narrow-spectrum growth inhibition was maximal with a cyclic cross-feeding network, while fully connected cross-feeding networks maximized coexistence during broad-spectrum growth inhibition. However, the effects of cyclic cross-feeding were particularly sensitive to community size and the number of targeted species; for communities with more than six species and resources, cyclic cross-feeding can be detrimental to coexistence due to instability. Our findings highlight the complex effects of cross-feeding network architecture on coexistence when growth inhibition reshapes the nutrient competition landscape, with potential applications to microbial communities in all natural environments, where they are often exposed to agents such as drugs, temperature, and pH modulation.

## Introduction

The gut microbiome is a dynamic ecosystem in which microbial species interact through various mechanisms, including nutrient competition, cross-feeding, and the secretion of inhibitory compounds. Perturbations such as drugs, temperature shifts, and dietary changes can disrupt this delicate balance by differentially altering bacterial cell growth and survival across the community, leading to shifts in microbiome composition and function. A case in point is antibiotics, which are administered to treat bacterial infections but inevitably also target the commensal bacteria that live in the gut^1,2^. Antibiotic treatments can reduce gut microbial diversity^3^, leaving the host more susceptible to infections of pathogens^4^. Successful prediction of the gut microbiome response to antibiotic treatments could enable a reduction in the rate of unintended consequences from antibiotics. However, our ability to predict treatment outcomes is currently limited by a lack of understanding of the interplay between antibiotics and community context^5^. Antibiotics induce cell death and/or reduce growth rate^6^, and are often studied using a single species in monoculture^7^. However, the community context of other species present in the environment can affect the susceptibility of a microbial species to antibiotic treatment^8^, due to the many mechanisms by which microbial species interact^3^. Such interactions include resource competition^9,10^ and cross-feeding (the secretion of metabolites that are consumed by other species)^11-14^, among others^15^.

Nutrient competition is a dominant factor in the assembly of human gut commensals into communities *in vitro* both without^16^ and with antibiotics^17^. Consumer-resource (CR) models in which species compete for steadily supplied resources have proven a useful tool for probing microbial community dynamics^18^ and predicting when species can coexist^19-21^. While CR models of chemostat-like growth typically assume that bacterial death rates are negligible relative to the dilution rate, this assumption breaks down during antibiotic treatment. In a recent study, we investigated the effects of incorporating species-specific death rates to a CR model, which highlighted the complex interplay between resource competition and antibiotic targeting regardless of the mechanism by which the antibiotic acts^22^. While mathematical models generally suggest that resource competition limits diversity (the competitive exclusion principle)^23^ and that species-specific death rates often reduce diversity further^22^, our study showed that antibiotics can actually promote diversity in a dose-dependent manner by inhibiting competitors that would win out otherwise^22^. However, this study ignored all other mechanisms of interspecies interactions besides nutrient competition, including cross-feeding.

In the absence of antibiotics or other growth-inhibiting stresses, cross-feeding can in principle increase community diversity^11,24^, with significant impact on the health of the host^25-27^. Cross-feeding interactions such as those in the human gut microbiome form a complex map^13,14,25,28^, hence theoretical studies of the interplay between competition and cooperation can be a useful tool to understand human gut microbiome population dynamics^29^. Cross-fed nutrients can be consumed predominantly by a single other species^30-33^, broadly by many other species^31,32^, or primarily by the species leaking the nutrient (selfish)^34,35^. The interplay among these distinct cross-feeding networks, nutrient competition, and antibiotic treatment have yet to be captured within a comprehensive modeling framework, despite the critical importance for accurately predicting the effects of antibiotics on gut microbiota diversity^30,36^.

Here, we develop a modified consumer-resource model with cross-feeding (resource leakage) to quantify how resource competition, cross-feeding network architecture, and stress-induced death or growth inhibition collectively determine stable coexistence. We consider selfish, cyclic, and fully connected cross-feeding networks, and define a leakage angle that parameterizes a spectrum spanning these network extremes. Across many leakage rates and extents of resource competition, the cross-feeding network that maximized the probability of maximum diversity (coexistence fraction) depended sensitively on the number of species and resources present (community size) and the number of species targeted by the growth-inhibiting stress (analogous to antibiotic spectrum). We found that cross-feeding can reduce diversity by introducing instability (dependent primarily on community size) or promote diversity by restructuring the nutrient competition landscape in favor of the targeted species (dependent primarily on treatment spectrum). We observed instability only for selfish and cyclic cross-feeding networks, and only for sufficiently large communities in the latter case. When the stress targeted a single species, coexistence fraction was highest for cyclic cross-feeding networks, while coexistence fraction was highest for fully connected networks when targeting multiple species, although the effect size of fully connected leakage largely decreased with community size.

## Results

### A CR model incorporating species-specific death rates and cross-feeding (leakage)

In a CR model of *m* microbial species competing for *p*steadily supplied resources in a well-mixed chemostat, the growth rate of species *i* is 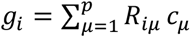 , where *R*_*iμ*_ is the rate at which species *i* consumes resource *μ* and *c*_*μ*_ is the concentration of resource *μ*. The rate of change of the abundance of species *i* is

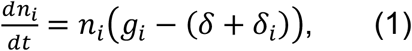

where *n*_*i*_ is the abundance of species *i, δ* is the dilution rate of the chemostat, and *δ*_*i*_ is the death rate of species *i*. The rate of change of resource *μ* is given by 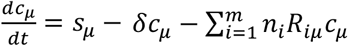 where *s*_*μ*_ is the supply rate of resource *μ*. The dilution term −*δc*_*μ*_ can be ignored for simplicity without significantly changing the behavior of the system^20^ (Fig. S1).

To incorporate cross-feeding in a general manner, we modified the CR model by assuming that a fraction of the mass of resource *v* consumed by species *i* is leaked as resource *μ* rather than contributing to the growth of species *i*. Since the CR model assumes that all resources are interchangeable in contributing to growth, we assume that the leakage fraction is independent of the identity of the consumed nutrient *v* and define *L*_*iμ*_ as the fraction of resource mass consumed by species *i* that is leaked as mass of resource *μ*. Hence, the fraction of total resource mass consumed by species *i* that contributes to the growth of species *i* is 1 − *L*_*i*_, where 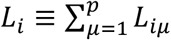 is the total leakage fraction of species *i*. This leakage modifies the growth rate of species *i* through both the factor 1 − *L*_*i*_ and the effect of leakage on the resource concentrations *c*_*μ*_:

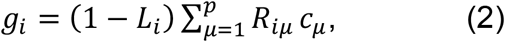

and the rate of change of the concentration of resource *μ* becomes

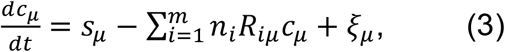

where ζ_*μ*_ is the total leakage of resource *μ*:

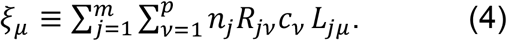

As in previous studies^20^, we can assume that resource consumption and leakage occur on much shorter time scales than cell division, leading to a separation of time scales such that *dc*_*μ*_/*dt* = 0. With this assumption, rearranging Eq. 3, the concentration of resource *μ* is 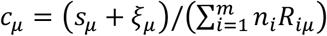. Plugging this solution into Eq. 2, Eq. 1 becomes

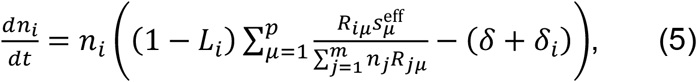

where we define the effective supply rate 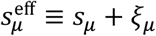 as the sum of the supply rate and the total leakage rate of resource *μ*. Eq. 1-5 collectively describe the CR model with leakage (CRL model) (Fig. 1A).

**Figure 1:**
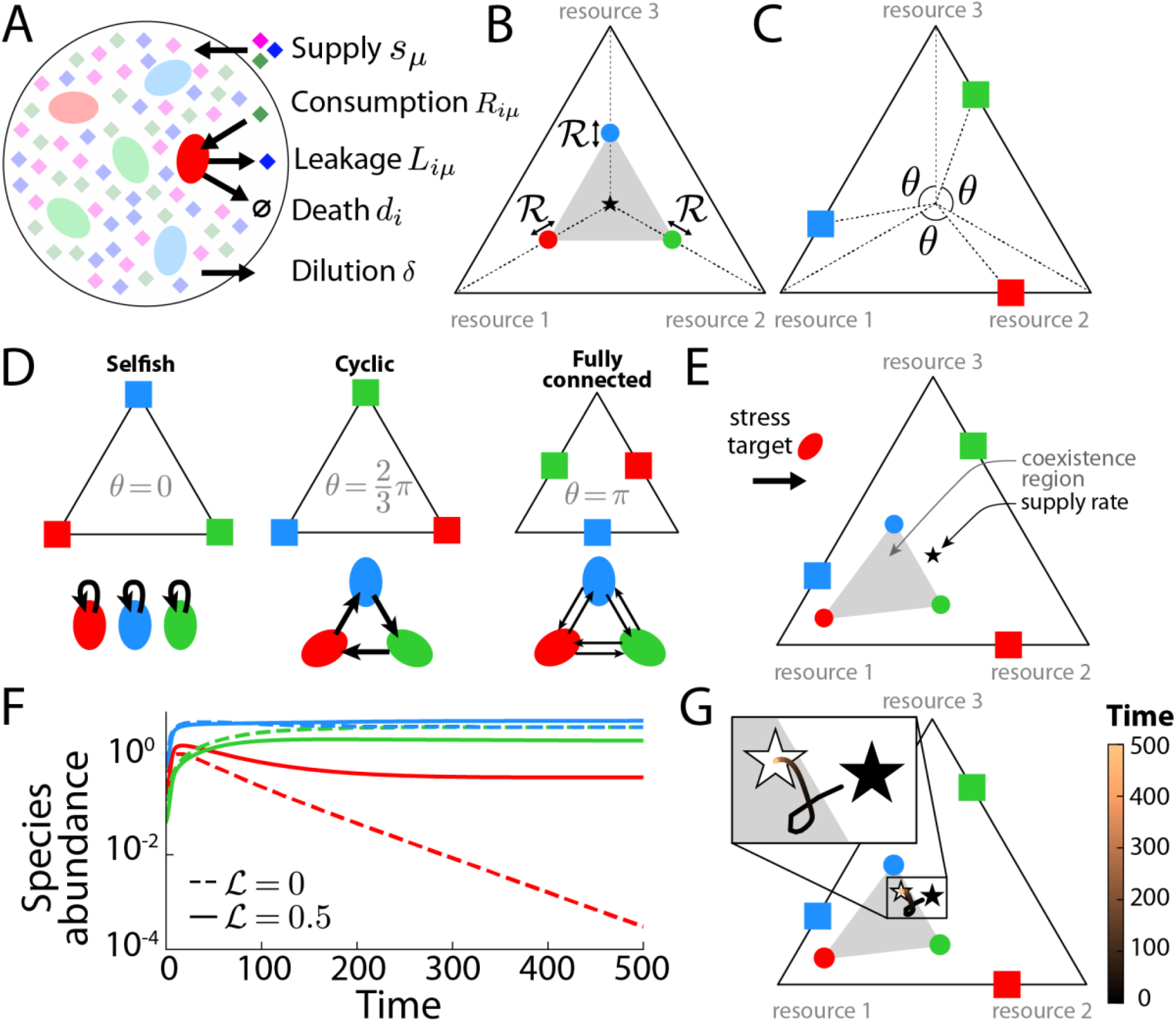
The consumer-resource model with species-specific death rates and resource leakage (CRL model). A) Three resources (diamonds) are supplied at rates *s*_*μ*_ to the chemostat in which three microbial species (ovals) consume resources, leak (cross-feed) resources, and die from a growth-inhibiting stress such as antibiotic treatment (or are diluted). B) Simplex depicting the relative resource consumption rates (circles), resource competition parameter ℛ, coexistence region formed by the convex hull of the resource consumption rates (gray region), and relative resource supply rates (solid star). C) Simplex depicting leakage angle *θ* and relative resource leakage rates (squares) (Methods). D) Simplex (top) and schematic (bottom) depicting selfish (*θ* = 0), cyclic (*θ* = 2*π*/3), and fully connected leakage (*θ* = *π*). E) When the death rate of the red species due to a growth-inhibiting stress is increased from zero, mimicking treatment with a narrow-spectrum antibiotic, the coexistence region is remapped toward the consumption niche of the red resource. F) Population dynamics without leakage (ℒ = 0, dashed lines) and with moderate total leakage fraction (ℒ = 0.5, solid lines) for the community in (E), showing that the red species goes extinct without leakage, but coexistence occurs with moderate leakage. G) The effective resource supply rate versus time for the moderate leakage simulation in (F), in which the color of the effective resource supply rate curve corresponds to the color bar in (F). The effective supply rate starts near the actual supply rate (solid star) and ends inside of the coexistence region, where the effective supply rate at the final time point of the simulation is shown by a white star.

By rescaling species abundances and resource consumption rates as 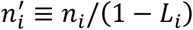 and 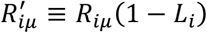, respectively, Eq. 5 becomes

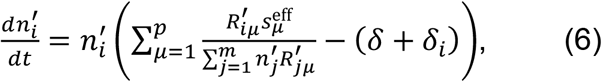

which is identical to the population dynamics equations in the CR model with species-specific death rates 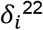^22^ up to rescaling species abundances 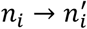 and resource consumption rates 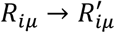 and shifting resource supply rates 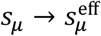.However, determining the effective resource supply rates 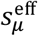 at steady state is non-trivial since the total microbial leakages ξ_*μ*_ (Eq. 4) are functions of the species abundances *n*_*i*_ and resource concentrations *c*_*μ*_. Furthermore, since the effective supply rates change during the approach to steady state, the dynamics and stability of communities in the CRL model are in general distinct from the corresponding community without leakage.

### A simplified case study of the effects of resource competition, cross-feeding, and death on species coexistence

As species can in principle invade the community until the number of coexisting species equals the number of supplied nutrients, we first assume the community saturates the bound set by the competitive exclusion principle (*m* = *p*). When *m* = *p* and the resource consumption rate vectors are linearly independent, we can calculate the unique steady-state resource concentrations and species abundances in the CRL model, assuming that all species abundances are non-zero (Methods). If the steady-state solution is physical (i.e., positive resource concentrations and species abundances), then we can calculate the Jacobian to determine its linear stability (Methods). We define coexistence as when the steady-state solution is physical and linearly stable (Methods).

We first consider communities of *m* = 3 species and *p* = 3 resources with total supply rate equal to unity 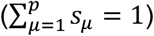, equal supply rates ( 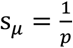 for all resources *μ*), and dilution rate *δ* = 0.1. We parameterize the resource consumption rate matrix *R* such that each species prefers a single, unique resource and has equal consumption rates of the other two non-preferred resources. That is, for species *i* and resource 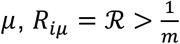 if *i* = *μ*, and *R*_*iμ*_ = (1 − ℛ)/(*m*− 1) if *i* ≠ *μ*; hence, the resource consumption rate vectors are linearly independent. Within this parameterization, we refer to resource *μ* as the niche of species *i* if *μ* = *i*. When visualized in a simplex (Fig. 1B), the convex hull of the resource consumption rates is an equilateral triangle, whereby larger triangles (larger ℛ) represent less resource competition. We relaxed the assumptions of uniform resource supply and symmetric resource consumption rates and found that our conclusions were largely unchanged (Fig. S2). In the absence of both growth inhibition and cross-feeding, a previous study showed that coexistence occurs if and only if the resource supply rates lie in the convex hull of the resource consumption rates^20^, so we refer to this region as the coexistence region (Fig. 1B, gray).

We focus on equal total leakage fractions (*L*_*i*_ = ℒ for all species *i*) and we visualize the leakage fractions *L*_*iμ*_ on the same simplex with the resource consumption rates (Fig. 1B). To systematically explore leakage scenarios, we represent leakage fractions as an equilateral triangle rotated by an angle *θ* ∈ [0, *π*], where *θ* = 0 is the case when species *i* leaks only its own niche resource (selfish leakage) (Fig. 1C, D). When *θ* = 2*π*/3, species 1, 2, and 3 leak the resource corresponding to the consumption niche of species 2, 3, and 1, respectively (cyclic leakage; Fig. 1D). When *θ* = *π*, each species leaks at equal rates the two resources corresponding to the consumption niches of the other two species (fully connected leakage; Fig. 1D). To account for a growth-inhibiting stress such as antibiotic treatment, we showed previously that when the death rate of a species is nonzero, the new coexistence region bounds can be calculated by determining the effective resource consumption rates (Fig. 1E)^22^.

Although selfish leakage may not appear beneficial in many biological contexts, molecules such as acetate are both consumed and secreted in some monocultures^35^. Moreover, selfish leakage could encompass the scenario in which a pair of species with very similar resource preferences assist each other and no other species through cross-feeding; in this case, the “species” engaging in selfish leakage would be the combined population of these mutualists. Finally, consumption of a resource in a given class may result in leakage of a closely related metabolite in that class to which the species in the community exhibit similar preferences as to the externally supplied resource^34^, which is similar in concept to selfish leakage. Indeed, our conclusions were largely unchanged if the leaked metabolites are distinct from the externally supplied resources but in the same metabolite class with regards to species preferences (Fig. S3).

We numerically integrated Eq. 1 and 3 to calculate the population dynamics of each combination of resource consumption parameter ℛ, death rates, and leakage fraction ℒ (Fig. 1F, G). For a community with intermediate ℛ and positive death rate for one species, mimicking treatment with a narrow-spectrum antibiotic, that species went extinct without leakage (Fig. 1F, dashed lines). However, coexistence occurred for intermediate total leakage fraction (ℒ = 0.5) and intermediate leakage angle *θ* (Fig. 1F, solid lines), due to the effective supply rates shifting to be inside the coexistence region as the system evolved away from an initial state with approximately equal abundances of all species (Fig. 1G). This example of a qualitative shift in coexistence motivated a systematic analysis of the impact of leakage network architecture.

### In the absence of death, small leakage angle and large leakage fraction can introduce instability, disfavoring coexistence

We first systematically tested the effect of leakage in the absence of death (*δ*_1_ = *δ*_2_ = *δ*_3_ = 0) and uniform supply of all three resources. We varied the remaining parameters independently in equally spaced intervals across their respective domains (ℛ ∈ [.35, .99], ℒ ∈ [0, 0.8], *θ* ∈ [0, *π*]), determining whether coexistence occurs (which we defined to be the existence of a physical, stable steady-state solution with all species abundances positive) for each (ℛ, ℒ, *θ*) (Methods). For each (*θ*, ℒ), we calculated the coexistence fraction, defined as the proportion of values of ℛ that resulted in coexistence (Fig. 2A). As expected, without leakage (ℒ = 0), the coexistence fraction was 1 since the uniform supply rates always lay in the coexistence region regardless of ℛ. For selfish leakage (*θ* = 0), increasing the total leakage fraction reduced the coexistence fraction due to the potential for a positive feedback loop whereby leakage of resource *i* increases the abundance of species *i*, which further increases leakage of resource *i*. This effect distinguishes selfish leakage from a scenario without leakage.

**Figure 2:**
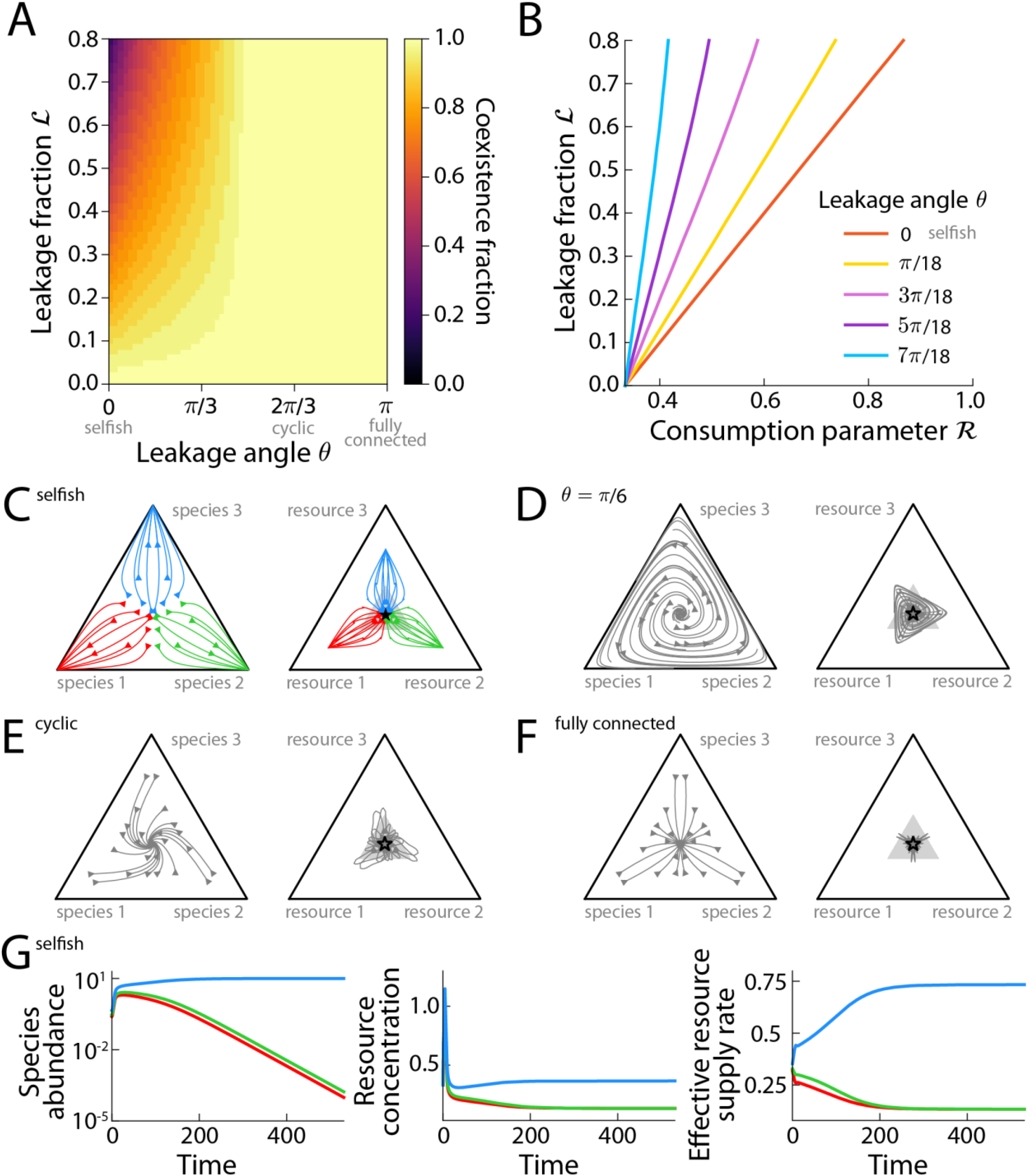
Without death, the coexistence fraction increases with leakage angle and decreases with total leakage fraction for sufficiently small leakage angles. A) The fraction of simulations that exhibited coexistence across variable extent of resource competition (ℛ ∈ [0.35, 0.99]) with zero death (𝒟 = 0) increased with leakage angle until *θ* reached *π*/2, after which coexistence occurred for all ℛ. B) For the simulations in (A), shown are the curves for five leakage angles below which coexistence occurred, and above which extinction occurred due to instability. C) For moderate leakage fraction (ℒ = 0.6), selfish leakage (*θ* = 0), and moderate resource competition (ℛ = 0.5), the initial relative species abundance impacted the final species abundance; the curves represent relative species abundances (left) and relative effective resource supply rates (right) over time, and the color of the curves depicts the species that persisted at abundance > 10^−2^ at the final time point of the simulation. For instance, initial abundances that were biased toward species 1 (closer to the lower left corner) biased the final state toward species 1 (red). D-F) Like (C), but for (D) *θ* = *π*/6, (E) *θ* = 2*π*/3 (cyclic leakage), and (F) *θ* = *π* (fully connected leakage). Gray curves indicate that all species persisted at abundance > 10^−2^ at the final time point of the simulation. G) Dynamics of species abundances, resource concentrations, and effective resource supply rates for the simulation in (C). The initial abundances of species 1, 2, and 3 were 0.25, 0.3, and 0.45, respectively, and the species with largest initial abundance (blue) outcompeted the other two species (green and red).

A similar effect occurred for small leakage angles 0 < *θ* < *π*/2 (Fig. 2A). However, for larger leakage angles representing cyclic-or fully connected-like leakage, an increase in the abundance of species *i* promoted coexistence by cross-feeding the other species, and coexistence always occurred for *θ* ≥ *π*/2 (Fig. 2A). We noticed that for each (*θ*, ℒ), coexistence occurred if and only if resource competition was low such that ℛ was greater than a critical consumption rate ℛ^*^(*θ*, ℒ) (Fig. 2B). Similarly, for each (*θ*, ℛ), coexistence occurred if and only if the total leakage fraction ℒ was smaller than a critical total leakage fraction ℒ^*^(*θ*, ℛ) (Fig. 2B). In other words, spontaneous symmetry breaking due to instability requires sufficiently large leakage fraction and sufficiently large extent of resource competition (small ℛ).

### Communities with cross-feeding exhibit multistability when extinction occurs

Experiments have shown that synthetic communities^17,37^ and human stool-derived *in vitro* communities^38^ can exhibit multiple stable steady states, such that the initial species abundances affect steady-state richness and/or abundances. While CR models involving only nutrient competition have a single steady state as long as all species are initially present^39^, the combination of cross-feeding and nutrient competition in the CRL model can lead to multistability (Fig. 2C,G), even when the consumption rates are not symmetric (Fig. S2), but communities with larger leakage angles often exhibited the same steady state across a wide range of initial abundances (Fig. 2D-F). To determine the regimes of multistability, we randomly sampled values of (ℛ, ℒ, *θ*) and calculated the population dynamics for three initial species abundance vectors with a low relative abundance of one or two species and determined whether the species abundances at long time scales were independent of the initial abundances (Fig. S4). When (ℛ, ℒ, *θ*) did not lead to coexistence, the steady state must be unstable, and hence any non-uniform initial species abundances will lead to extinction of at least one of the three species. In such a scenario, symmetry implies multiple steady states each with a distinct set of extinct species (multistability). Conversely, when a given (ℛ, ℒ, *θ*) led to coexistence, simulations showed that 0% (0/100) of communities were multistable. Hence, multistability likely occurs if and only if the community does not exhibit coexistence.

### During treatment with a narrow-spectrum stress, coexistence is most likely with cyclic leakage networks

We next explored how the interplay between resource competition and leakage affects coexistence when one of the species has a non-zero death rate, mimicking treatment with a narrow-spectrum antibiotic. We varied each parameter independently and in equally spaced intervals across their respective domains (ℛ ∈ [.35, .99], ℒ ∈ [0, 0.8], *θ* ∈ [0, *π*], *δ*_1_ = 𝒟 ∈ [0.01, 0.2]) while keeping the other death rates zero (*δ*_2_ = *δ*_3_ = 0), and determined when coexistence occurs. For each (*θ*, ℒ), we calculated the coexistence fraction across (ℛ, 𝒟). For all positive total leakage fractions ℒ > 0, we found that the coexistence fraction was largest for cyclic leakage (*θ* = 2*π*/3) and smallest for selfish leakage (*θ* = 0, Fig. 3A).

**Figure 3:**
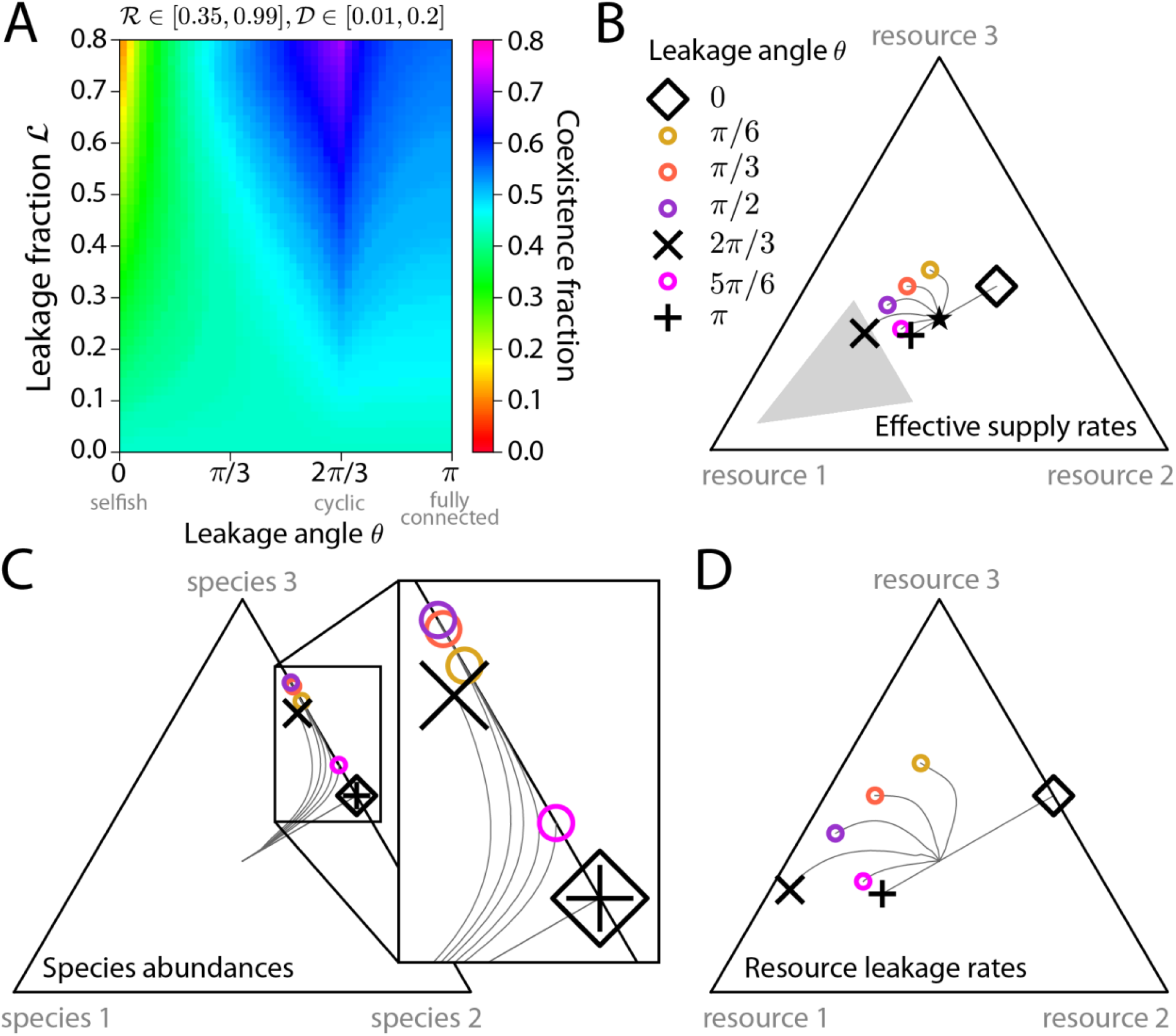
During a narrow-spectrum stress, coexistence is most likely with cyclic leakage. A) The coexistence fraction across death rates 𝒟 ∈ [0.01, 0.2] and extent of resource competition ℛ ∈ [0.35, 0.99] was approximately independent of leakage angle for small total leakage fraction (ℒ ≤ 0.2), but as the total leakage fraction increased above 0.2, the coexistence fraction increased (decreased) with ℒ for moderate and large *θ* (small *θ*). B-D) Dynamics of the relative effective supply rates (B), species abundances (C), and resource leakage rates (D) for various leakage angles, including selfish (*θ* = 0; black diamonds), cyclic (*θ* = 2*π*/3; black ×’s), and fully connected (*θ* = *π*; black +’s) leakage demonstrate that coexistence is most likely for cyclic leakage during a narrow-spectrum stress. Gray curves represent the dynamics and shapes are at the final time point, with (ℛ, ℒ, 𝒟) = (0.6,0.5,0.075).

To understand why communities with cyclic leakage were most likely to exhibit coexistence during a narrow-spectrum stress, we performed simulations with all species initialized at uniform abundances for (ℛ, ℒ, 𝒟) = (0.6,0.5,0.075) and compared population dynamics for seven leakage angles from 0 to *π*, including selfish (*θ* = 0, Fig. 3B-D, black diamonds), cyclic (*θ* = 2*π*/3, Fig. 3B-D, black ×’s), and fully connected (*θ* = *π*, Fig. 3B-D, black +’s) leakage. Regardless of leakage angle, the coexistence region shifted toward resource 1 (Fig. 3B) since species 1 was targeted with the stress. For all leakage angles, species 1 had the lowest abundance at steady state (Fig. 3C). For any leakage angle 0 < *θ* < *π*, species 1 leaks more of resource 2 than resource 3, thus inhibition of species 1 lowers the levels of resource 2 more than resource 3 and promotes species 3 over species 2 (Fig. 3C). Therefore, for any leakage angle 0 < *θ* < *π*, the majority of steady-state leakage is from species 3, so the relative leakage of resource 1 is maximized when species 3 exclusively leaks resource 1 (Fig. 3D; cyclic leakage). For these parameters, the effective supplied resource rates lay in the convex hull of the remapped resource consumption rates only for cyclic leakage (Fig. 3B), so this scenario was the only one in which coexistence occurred (Fig. 3C).

Thus, cyclic leakage in which species 3 accounts for >50% of the community at steady state (Fig. 3C) and solely leaks resource 1 is most beneficial for coexistence (Fig. 3B,C). By contrast, for fully connected leakage, even though both species 2 and 3 leak resource 1, the levels of relative resource 1 leakage are maximally 50%, when species 1 is extinct (Fig. 3D; black +’s), which is insufficient to rescue species 1 (Fig. 3C). For selfish leakage, the targeted species does not benefit from leakage by other species (Fig. 3B,D), so the coexistence fraction is lower than for nonzero leakage angle.

To study how the strength of the stress (e.g., antibiotic dose) affects these findings, we computed the coexistence fraction across a range of ℛ with fixed death rate 𝒟 = 0.01 (low compared to dilution rate), 0.1 (moderate; equal to dilution rate), or 0.2 (large antibiotic dose). For each 𝒟, the coexistence fraction was maximized for cyclic leakage (Fig. S5). For some leakage angles between selfish and cyclic leakage (e.g., *θ* = *π*/4), increasing the leakage fraction with 𝒟 = 0.01 resulted in non-monotonic richness changes (Fig. S6). Analogous to our previous work showing that changing the death rate can cause a community to transit through multiple coexistence states separated by extinction of different species^22^, we found that changing the leakage fraction affected which species go extinct. With ℛ = 0.5, as the total leakage fraction ℒ was increased from zero, coexistence was eliminated due to extinction of the targeted species (species 1, Fig. S6). Further increase in ℒ resulted in a switch to coexistence, and then extinction of the non-targeted species 2 due to the cyclic component of leakage at angle *θ* = *π*/4 (Fig. S6). This result contrasts with the case without death (Fig. 2B), when coexistence occurred if and only if the leakage fraction was below a critical value.

In summary, during narrow-spectrum stress treatment, our CRL model predicts that coexistence fraction will decrease with increasing total leakage fraction for small leakage angle (e.g., selfish, *θ* = 0), increase for large total leakage angle (e.g., cyclic or fully connected, *θ* = 2*π*/3 or *π*, respectively), and depend non-monotonically on total leakage fraction for intermediate leakage angle.

### During treatment with a broad-spectrum stress, coexistence is most likely with fully connected leakage networks

We next explored how leakage scenario affects coexistence when two of the three species have a non-zero death rate, mimicking treatment with a broad-spectrum antibiotic. We varied the parameters independently and in equally spaced intervals across their respective domains (ℛ ∈ [.35, .99], ℒ ∈ [0, 0.8], *θ* ∈ [0, *π*], *δ*_1_ = *δ*_2_ = 𝒟 ∈ [0.01, 0.2]) with *δ*_3_ = 0, and determined when coexistence occurred. For each (*θ*, ℒ), we calculated the coexistence fraction across (ℛ, 𝒟). For all positive total leakage fractions ℒ > 0, we found that the coexistence fraction was largest for fully connected leakage (*θ* = *π*, Fig. 4A), in contrast to a narrow-spectrum stress (Fig. 3A). Moreover, the coexistence fraction was non-monotonic with leakage angle, as the coexistence fraction was reduced when the leakage angle was perturbed slightly from the cyclic network (Fig. 4A), as with a narrow-spectrum stress (Fig. 3A).

**Figure 4:**
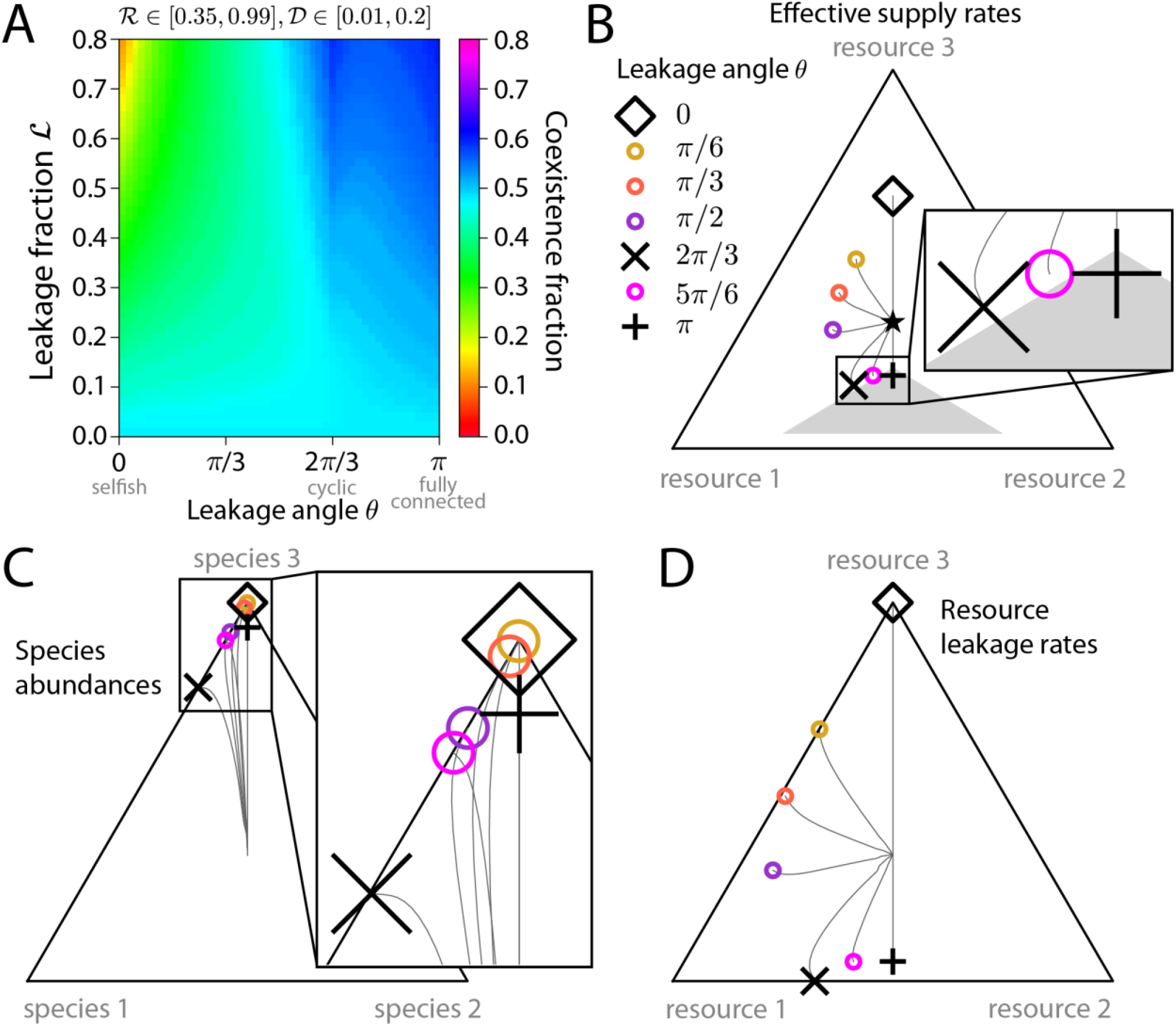
During a broad-spectrum stress, coexistence is most likely with fully connected leakage. A) The coexistence fraction across death rates 𝒟 ∈ [0.01, 0.2] and extent of resource competition ℛ ∈ [0.35, 0.99] was approximately independent of leakage angle for small total leakage fraction (ℒ ≤ 0.1), but as the total leakage fraction increased above 0.1, the coexistence fraction increased most for large leakage angles (i.e., fully connected leakage). B-D) Dynamics of the relative effective supply rates (B), species abundances (C), and resource leakage rates (D) for various leakage angles, including selfish (*θ* = 0; black diamond), cyclic (*θ* = 2*π*/3; black x), and fully connected (*θ* = *π*; black +) leakage, demonstrate that coexistence is most likely for fully connected leakage during a broad-spectrum stress. Gray curves represent the dynamics and shapes are at the final time point, with (ℛ, ℒ, 𝒟) = (0.6,0.5,0.065).

To understand why communities with fully connected leakage were most likely to exhibit coexistence during treatment with a broad-spectrum stress, we performed simulations in which all species were initialized at uniform abundances for (ℛ, ℒ, 𝒟) = (0.6,0.5,0.065) and compared the dynamics for various leakage angles, including selfish (*θ* = 0), cyclic (*θ* = 2*π*/3), and fully connected (*θ* = *π*) leakage. Regardless of leakage angle, the coexistence region shifted away from resource 3 (Fig. 4B) since species 1 and 2 were targeted with death, in contrast to the shift under a narrow-spectrum stress (Fig. 3B). For all leakage angles, species 3 was the most abundant at steady state (Fig. 4C). For fully connected leakage (*θ* = *π*), species 3 leaks resources 1 and 2 at equal rates such that there was sufficient relative steady-state leakage of resources 1 and 2 (Fig. 4D) to promote species 1 and 2 (Fig. 4B). For any leakage angle 0 < *θ* < *π*, species 3 leaks more of resource 1 than resource 2 (Fig. 4D), thereby increasing the levels of resource 1 more than resource 2 (Fig. 4B) and promoting species 1 over species 2 (Fig. 4C). As a result, species 2 went extinct for 0 < *θ* < *π* (Fig. 4C). As with a narrow-spectrum stress (Fig. 3B-D), communities with selfish leakage (Fig. 4B-D) did not exhibit coexistence during a broad-spectrum stress as both targeted species went extinct.

For cyclic leakage (*θ* = 2*π*/3), the relative steady-state leakage of resource 3 was minimized (Fig. 4D) since resource 3 is solely leaked by species 2, the least abundant species (Fig. 4C). Furthermore, the relative steady-state leakage of resource 2 increased with leakage angle (Fig. 4B). Coexistence is promoted by both minimizing the relative leakage of resource 3, which occurs at *θ* = 2*π*/3, and by maximizing the relative leakage of resource 2, which occurs at *θ* = *π*; these two effects underlie the non-monotonicity of coexistence fraction with increasing leakage angle for large total leakage fraction (Fig. 4A).

We explored how the strength of a broad-spectrum stress affects our findings by computing the coexistence fraction across a range of ℛ with fixed 𝒟 = 0.01, 0.1, or 0.2. As with a narrow-spectrum stress, we found that the coexistence fraction was maximized for the same leakage network (fully connected in this case) regardless of dose (Fig. S7). Thus, the interplay between resource competition and leakage network architecture can substantially affect microbial community coexistence during growth inhibiting treatments, with qualitatively distinct outcomes depending on how many community members are affected by the antibiotic.

### With cyclic leakage, coexistence fraction depends nonmonotonically on community size during growth inhibition

We next queried how the findings above for small communities (3 species and 3 resources) generalize to larger communities. First, we quantified coexistence outcomes during a narrow-spectrum stress with cyclic or fully connected leakage as we increased the community size by increasing the starting number of species *m* and number of resources *p* such that *m* = *p*. To do so, we generalized our definition of the two leakage network architectures: for fully connected leakage, *L*_*iμ*_ = 0 if *i* = *μ*, and *L*_*iμ*_ = *L*_*i*_/(*m*− 1) if *i* ≠ *μ*; for cyclic leakage, *L*_*iμ*_ = *L*_*i*_ if *i* + 1 ≡ *μ* (mod *m*), and *L*_*iμ*_ = 0 otherwise. We defined a narrow-spectrum stress as one that results in a nonzero death rate for only one species 1 (*δ*_1_ = 𝒟, *δ*_*i*_ = 0 for *i* > 1). Like in the previous sections, we studied symmetric resource consumption rate matrices parametrized by ℛ. We uniformly sampled the parameters ℛ ∈ [.35, .99], ℒ ∈ [0, 0.8], 𝒟 ∈ [0, 0.2], *m* = *p*∈ [3, 40], leakage network *N*_*L*_ ∈ {none, cyclic, fully connected}, and determined whether coexistence occurs for each (ℛ, ℒ, 𝒟, *m, N*_*L*_). We then calculated the coexistence fraction for subsets of these parameters (Fig. 5A).

**Figure 5:**
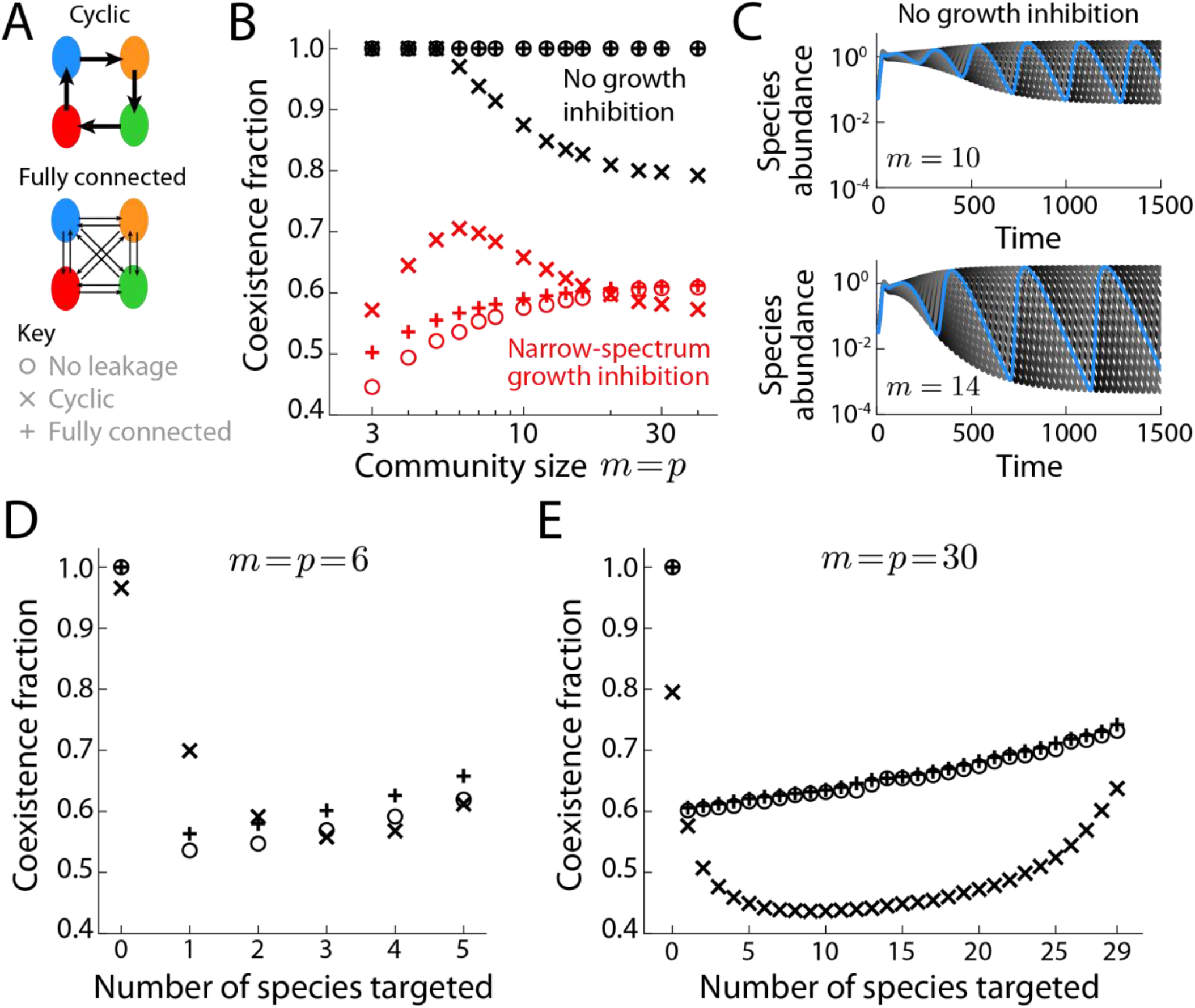
The coexistence fraction for growth-inhibited communities varies with community size, leakage network architecture, and targeting spectrum. A) Schematic depicting cyclic (top) and fully connected (middle) leakage for a community with *m* = *p* = 4, and a key for leakage labels in (B,D,E). B) Coexistence fraction across extent of resource competition ℛ ∈ [0.35, 0.99], total leakage fraction ℒ ∈ [0.1, 0.8], and death rate of targeted species 𝒟 ∈ [0, 0.2] for fully connected (+’s), cyclic (×’s), and no leakage (o’s), and without death (black) or during a narrow-spectrum stress in which one species is targeted (red). The coexistence fraction increased with community size for fully connected and selfish leakage, and was maximal at intermediate community size for cyclic leakage. C) Species abundance dynamics for communities of *m* = 10 (top) and *m* = 14 (bottom) species with cyclic leakage and (ℛ, ℒ) = (0.5, 0.8), in the absence of death. Simulations were initialized with one species at lower abundance (blue) and all others at the same, larger abundance (gray curves). Both communities were unstable; for the larger community (bottom), the period and amplitude of oscillations in the abundance of each species (highlighted by the blue curve) was larger than for the smaller community (top). D) For a community with *m* = *p* = 6, the coexistence fraction varies with the number of species targeted. For fully connected (+’s) and cyclic (×’s) leakage, each point is a mean across (ℛ, ℒ, 𝒟) values, and for no leakage (o’s), each point is a mean across variable (ℛ, 𝒟) values, with death rate 𝒟 ∈ [0.01, 0.2], extent of resource competition ℛ ∈ [0.35, 0.99], and total leakage fraction ℒ ∈ [0.1, 0.8]. During a broad-spectrum stress, fully connected leakage resulted in higher coexistence fraction than with cyclic leakage or without leakage. Note that the scenario in which all six species are targeted is identical to targeting zero species. E) Like (D), but for a larger community (*m* = *p* = 30). For this community, cyclic leakage is detrimental to coexistence and its effects depend strongly on the number of targeted species, while the effects of fully connected leakage on coexistence are negligible regardless of targeting spectrum.

Without death (𝒟 = 0), the coexistence region is defined by the resource consumption rate vectors. In the absence of leakage, a supplied resource rate vector within the convex hull leads to coexistence^20^. Since fully connected and cyclic leakage do not inherently favor any particular resource, these mechanisms only cause extinction if they result in spontaneous symmetry breaking, as we observed for selfish leakage (Fig. 2). We found that for fully connected leakage, the coexistence fraction was 1 for all community sizes (Fig. 5A, black +’s), but for cyclic leakage, the coexistence fraction decreased from 1 as the community size increased above *m* = *p* = 6 (Fig. 5A, black ×’s), indicating that symmetric breaking can occur for some (ℛ, ℒ). Indeed, the steady state was unstable for large total leakage fraction ℒ and high resource competition (low ℛ) (Fig. 5B), unlike for *m* = *p* = 3 (Fig. 2D). For ℛ = 0.5, ℒ = 0.7, and *m* = *p*> 6, species abundances exhibited oscillations, in which any decrease of species *i* initially reduces the leakage of resource *i* + 1 (mod *m*) and thus reduces the abundance of species *i* + 1 (mod *m*). However, these decreases in species abundance free up more resources, which allows species *i* to eventually recover, thereby providing more resources for species *i* + 1 (mod *m*) and driving its subsequent recovery (Fig. 5B).

Intuitively, increasing ℒ or increasing the extent of resource competition by reducing ℛ should increase the extent to which a small drop in abundance of species *i* reduces the abundance of species *i* + 1 (mod *m*), consistent with our numerical findings (Fig. 5A). The amplitude of the oscillations increased over time, and the period and rate of amplitude growth increased with increasing community size (Fig. 5B). In the absence of leakage, treatment with a narrow-spectrum stress (Fig. 5A, red o’s) reduced the coexistence fraction relative to the absence of death (Fig. 5A, black o’s), as expected due to growth inhibition of the targeted species. However, the coexistence fraction increased with community size (Fig. 5A, red o’s), consistent with our previous study^22^ in which we found that adding an additional species and resource shifts the effective coexistence region and supply rate such that the net effect is to reduce competition for the targeted species’s resource niche (Fig. S8A).

Introducing fully connected leakage to a community treated with a narrow-spectrum stress (Fig. 5A, red +’s) increased the coexistence fraction regardless of community size, but the effect was negligible for large community sizes. Fully connected leakage promotes coexistence since all non-targeted species leak the resource preferred by the targeted species, but this effect is dampened as community size increases since the targeted species becomes more of a generalist consumer (Fig. S8A) and hence the fully connected cross-feeding is less beneficial.

Introducing cyclic leakage to a community treated with a narrow-spectrum stress (Fig. 5A, red ×’s) increased the coexistence fraction for small and intermediate community sizes (*m* = *p* ≲ 20) but not for larger communities (*m* = *p* ≳ 20). For this leakage network, the effect was largest for intermediate community size, and the coexistence fraction was maximal at *m* = *p* = 6. For small community sizes (*m* = *p*< 6), cyclic leakage promotes coexistence via a non-targeted species leaking the resource preferred by the targeted species without leading to instability. However, for larger community size (*m* = *p*> 6), the instability induced by cyclic leakage (Fig. 5B) decreases the coexistence fraction. Thus, both fully connected and cyclic leakage increase the coexistence fraction during a narrow-spectrum stress for small and intermediate community sizes (*m* = *p* ≲ 20); for larger communities (*m* = *p* ≳ 20), fully connected leakage has a negligible effect on coexistence fraction, and cyclic leakage reduces the coexistence fraction relative to the absence of leakage (Fig. 5A, red).

To investigate broad-spectrum stresses, we focused on a community of intermediate (*m* = *p* = 6) and large (*m* = *p* = 30) size. For each community size, we compared the coexistence fraction for fully connected, cyclic, and no leakage as we varied the number of species targeted with a death rate 𝒟 (with zero death rate for the the other non-targeted species). For *m* = *p* = 6, when at least one species was targeted, without leakage the coexistence fraction increased with the number of targeted species because of resource competition (Fig. 5C, o’s), since there were fewer non-targeted species that could outcompete the targeted species for nutrients and drive extinction (Fig. S8B). Like a narrow-spectrum stress (Fig. 5A), fully connected leakage (Fig. 5C, +’s) resulted in a small increase in coexistence fraction relative to no leakage as long as at least one species was targeted. In contrast, cyclic leakage resulted in a small decrease in coexistence fraction relative to no leakage if > 2 species were targeted, despite this network having the highest coexistence fraction for a narrow-spectrum stress (Fig. 5A). The coexistence fraction for cyclic leakage was non-monotonic, with a minimum at *n* = 3 targeted species (Fig. 5C, ×’s). These findings were largely conserved for a larger community with *m* = *p* = 30 (Fig. 5D), with slightly lower coexistence fractions for all number of targeted species with cyclic leakage, consistent with the increased likelihood of instability in large cyclic communities (Fig. 5B). As with *m* = *p* = 6, the coexistence fraction of cyclic communities was minimal when an intermediate number of the species were targeted (Fig. 5D, ×’s).

Taken together, our findings from communities with three members were generally applicable to communities of up to six species. However, for large communities with >20 species, cyclic leakage is detrimental to coexistence in the absence or presence of growth-inhibiting stresses and its effects depend strongly on the number of targeted species, while the effects of fully connected leakage are negligible regardless of targeting spectrum.

## Discussion

To simplify our analyses, we focused on scenarios in which each species in the community was an equal contributor to the same leakage network. Nonetheless, our findings suggest a complex interplay between leakage, resource competition, and death; for instance, in certain regimes, leakage angles corresponding to cyclic and fully connected networks are more likely to result in coexistence than leakage angles in between (Fig. 4A). Our findings suggest cross-feeding is a benefit to the diversity of small communities during growth-inhibiting stresses such as antibiotic treatments if the cross-feeding network is not primarily selfish (Fig. 3,4). In the absence of leakage, we previously found that the qualitative effects of such stresses were largely insensitive to community size. In contrast, our CRL model predicts that large networks of cyclic cross-feeding species are more likely to exhibit instabilities (Fig. 5B), suggesting a selective pressure against such networks, and that the consequences of cross-feeding can either be amplified or dampened (Fig. 5A,C) as cross-feeding networks expand in size. In general, our results suggest that the interplay between growth-inhibiting stresses and cross-feeding is likely to be most evident in smaller communities, which tend to be more tractable experimentally^36^. Future work could consider more general forms of the leakage fraction matrix; for example, in a three-species, three-resource community, the leakage rates could lie anywhere in the simplex.

Experimental validation of CR models will rely on the ability to quantify and tune consumption and production rates within microbial communities. These rates can be parameterized through growth measurements coupled to metabolomics profiling^40^ and tuned using CRISPRi knockdown^41^ if the organisms are genetically tractable. Although cross-feeding can play important roles in natural bacterial communities^11-14^, quantitative knowledge about cross-feeding interactions that are relevant for coexistence is currently limited. Testing the predictions of our CRL model will require the establishment of multiple communities with distinct cross-feeding network architectures, parameterized consumption and leakage rates, and leakage substantial enough to impact coexistence during stresses, as well as the ability to tune leakage to establish that the community exhibits coexistence in the absence of leakage or stress. For microbial communities in nature, strains/species are likely to exhibit a wide range of leakage magnitudes among many subnetworks, potentially with different architectures^42^. To parameterize such communities, it will be useful to establish general relationships between cross-feeding and other physiological variables, such as whether the magnitude of leakage is linked to overall growth rate. A community such as the gut microbiota with hundreds of species^43^ may contain only a few that engage in leakage, while many other members focus on resource competition^16^. In these cases, even if there are multiple subnetworks, our results for simplified communities could be applied to understand the coexistence of a subset of the larger community under a growth-inhibiting stress treatment, especially if there is a dominant subnetwork with the largest effect on community assembly. Moreover, the framework we have developed could be used in the future to interrogate coexistence of communities that exhibit a combination of multiple leakage subnetworks. Using block-diagonal leakage matrices in which each block corresponds to a sub-community of species that interact through cross-feeding, it may be possible to determine to what extent a community with multiple subnetworks behaves as the superposition of the single-network communities studied here.

Building on previous formulations of cross-feeding^18,20,34,44-49^, which generally focus on the interplay between resource competition and leakage, our framework naturally defines a spectrum of cross-feeding networks and the interplay with the spectrum of stress targeting in the context of classical consumer-resource models. The network(s) present within a community could be quantified experimentally through time-resolved metabolomics measurements of resource leakage dynamics. Such measurements would provide information about the size of cross-feeding networks, and guide experiments focused on mutants lacking cross-feeding capacity that are most likely to disrupt coexistence during a growth-inhibiting stress. While persistent fluctuations^50^, instability^29^, chaos^51^, and multistability^18,52,53^ in microbial communities have been studied theoretically and experimentally, the extent to which the interplay between cross-feeding and stresses generates multistability has remained largely unexamined; experimental tracking of e.g., antibiotic treatment coupled with computational modeling may help to identify such possibilities. Although antibiotic treatment is a natural context for the results in this study, our model is equally applicable to other factors that affect bacterial growth, such as pH and temperature changes (which broadly affect all species in the community). Our model can also be extended to address the effects of toxins^42,54^, which are effectively cross-fed compounds with a negative impact on growth, by incorporating death rates dependent on toxin concentrations. More generally, our findings indicate that extending consumer-resource models to incorporate ecologically relevant interactions beyond cross-feeding and resource competition may be necessary to fully understand and engineer microbial community dynamics.

## METHODS

### Closed-form steady-state solution to population dynamics equations

When the number of resources and species are equal (*m* = *p*), one can solve for the unique steady-state solution of species abundances and resource concentrations. To do so, we first rewrite Eq. 1-4 in matrix notation involving the species abundance vector 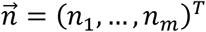 and the species death rate vector 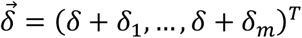 in which we incorporate the dilution rate *δ* into each component. Eq. 1 becomes

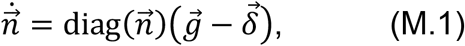

where 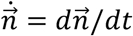 is the diagonal matrix with the elements of 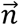 on the diagonal, and the growth rate vector is

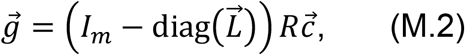

where *I*_*m*_is the *m*× *m*identity matrix, 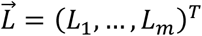 is the vector of leakage fractions, *R* is the matrix of resource consumption rates *R*_*iμ*_ , and 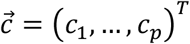 is the vector of resource concentrations. The rate of change of resource concentrations is

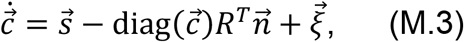

where 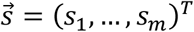 is the resource supply rate vector and 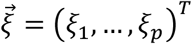 is the total microbial leakage of resources. Rewriting Eq. 4 in matrix notation,

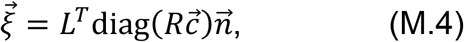

where *L* is the matrix of leakage rates *L*_*iμ*_.

To calculate the steady-state species abundances and resource concentrations, we set 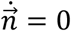 and assume all steady-state species abundances are non-zero. With this assumption, Eq. M.1 implies that 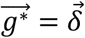, where the asterisk signifies the steady-state value. Plugging this relation into Eq. M.2 yields the steady-state resource concentrations

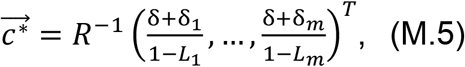

where we have utilized the fact that *R* is invertible for all systems in this study. Since 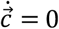 at steady state, Eq. M.3 and M.4 imply that 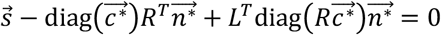 We can factor out 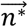 and rearrange to obtain

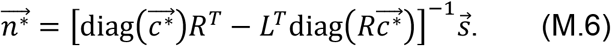

### Linearization of dynamical equations near the fixed point to determine stability

If 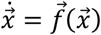, then the steady-state solution 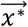 satisfies 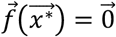 The Jacobian is defined as

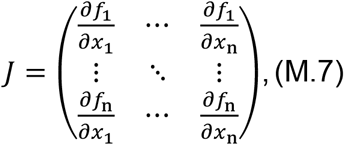

and the solution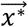 is stable if all eigenvalues of *J* at 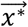 have negative real parts and unstable if any eigenvalue has a positive real part.

To determine whether the steady-state solution given by Eq. M.5 and M.6 is stable, we calculate the Jacobian matrix *J*. We write the CRL model equations in the form of Eq. M.7 by defining

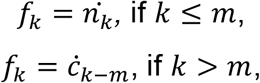

and

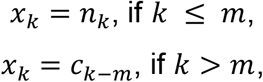

where *k* ∈ {1, … , *m*+ *p*}, and *m* = *p*. Next, we explicitly calculate the partial derivatives and evaluate them at steady state:

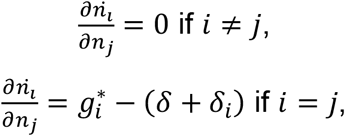

where 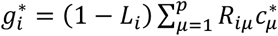 and

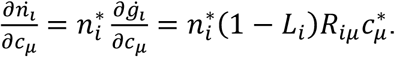

Moreover,

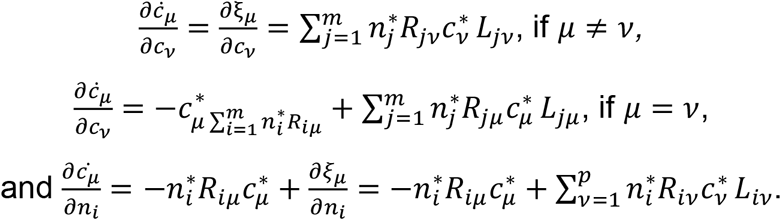

We numerically calculate the eigenvalues of the Jacobian matrix with the elements above. In all calculations for this study, all eigenvalues had non-zero real part.

### Simulations of population dynamics

We simulated species abundances over time in Python using the SciPy function scipy.integrate.solve_ivp, an explicit Runge-Kutta method for solving ordinary differential equations. Except for Fig. S3E, we integrated all species abundances *n*_*i*_ from time *t* = 0 to *t* = 10,000; in Fig. S3E, we integrated up to *t* = 4,000. In a small time step Δ*t*, the mass of species *i* that is removed due to dilution is *n*_*i*_(*t*)*δ*Δ*t*. Assuming that the abundance of species *i* is constant over time, if the dilution time scale of the chemostat is *δ* = 0.1, then 1000 times the mass of species *i* will be removed over the course of the simulation.

### Definition of coexistence and extinction

For a model microbial community defined by the parameters in Table 1, we sought to determine whether all species would stably coexist. We used two independent approaches: an analytic approach in which we calculated the fixed points of the dynamical equations and determined linear stability near the fixed points as described in the methods sections above, and numerical simulations.

**Table 1:** Description of all model parameters and their default values.

| Label | Definition |
| --- | --- |
| $m$ | Number of species. |
| $p$ | Number of resources. In this study, we considered systems with an equal number of species and resources ($m = p$). |
| $n_i(t = 0)$ | Initial abundance of species $i$. |
| $c_\mu(t = 0)$ | Initial concentration of resource $\mu$. In simulations, we set the initial resource concentrations to be uniform and sum to unity. |
| $R_{i\mu}$ | Rate at which species $i$ consumes resource $\mu$, per unit of species abundance per unit of resource concentration. We parameterize the matrix $R_{i\mu}$ by the extent of resource competition parameter $\mathcal{R}$. |
| $L_{i\mu}$ | Leakage fraction of resource $\mu$ by species $i$. In a given time interval, if species $i$ consumes a total resource amount $X$, then it leaks $XL_{i\mu}$ of resource $\mu$. In all cases, we set the total leakage fraction for species $i$ $L_i = \sum_\mu L_{i\mu}$ to a constant $L_i = \mathcal{L}$ for all species $i$. When $m = p = 3$, we parameterize $L_{i\mu}$ by the rotation angle $\theta$ in the unit simplex; for larger communities ($m = p > 3$) we considered two cases for nonzero $L_{i\mu}$ (fully connected and cyclic). |
| $\delta_i$ | Death rate of species $i$. |
| $\delta$ | Dilution rate of the chemostat ($\delta = 0.1$). |
| $s_\mu$ | Supply rate of resource $\mu$. In simulations, we set the supply rate to be uniform across all resources, sum to unity, and remain constant over time. |

For the analytical approach, we ignored fixed points for any species with a steady-state abundance of zero since this value corresponds to extinction, and we found that there is a unique fixed point for all parameter sets such that all species abundances are non-zero. If this unique fixed point is physical (i.e., all abundances are positive) and linearly stable, then we define the community to exhibit coexistence. This conclusion implicitly assumes that, for all initial conditions, the system will approach the fixed point with all positive species abundances rather than a fixed point with some species abundances zero. Our non-exhaustive testing of this assumption suggested that it is valid (Fig. 2E,F), although we did not prove it mathematically.

In simulations, if the abundance of a species dropped below 10^-4^, it was set to zero, signifying extinction and removal from the community, and we stopped the simulation at that time rather than continuing to *t* = 10,000. If all species abundances were greater than 10^-4^ at the final time step, we determined that coexistence occurred. This definition of coexistence is not necessarily equivalent to the analytical approach described above, although in practice we found that the two typically overlapped. Unless otherwise stated, we focused on the analytical approach, and used simulations to supplement our conclusions.

### Calculating coexistence fraction across parameter sets

When calculating the coexistence fraction across values of (𝒟, ℛ, ℒ, *θ*), we fixed a subset of parameters and varied the remaining parameters independently in equally spaced intervals across their respective domains, determining whether coexistence occurred for each instance of the parameter values.

### Visualization of parameters in simplex

We visualized the resource supply rates 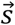 resource consumption rates *R*_*ij*_, and resource leakage fractions *L*_*ij*_ on a single simplex (Fig. 1B). We calculated the position of the resource supply rates (*s*_1_, *s*_2_, *s*_3_) by normalizing them to sum to unity (*s*_1_ + *s*_2_ + *s*_3_ = 1) and then plotting the linear combination *s*_1_**e**_1_ + *s*_2_**e**_2_ + *s*_3_**e**_3_, where **e**_*i*_ are the vectors representing each corner of the simplex. We similarly plotted the resource consumption rates (*R*_*i*1_, *R*_*i*2_, *R*_*i*3_) and relative leakage fractions (*L*_*i*1_, *L*_*i*2_, *L*_*i*3_) for species *i* on the same simplex.

### Without leakage, larger communities behave like communities of two species and two resources

As shown in our previous study^22^, in the absence of cross-feeding (Fig. 5A,C,D, o’s), the steady-state behavior of a community can be mapped onto a corresponding two-species, two-resource community in which one species is targeted with the stress and the other species has zero death rate (Fig. S8). As an example, consider a community with *m* = *p* = 5, in which three species have death rate 𝒟 and the other two species have zero death rate. The resource consumption rate matrix is

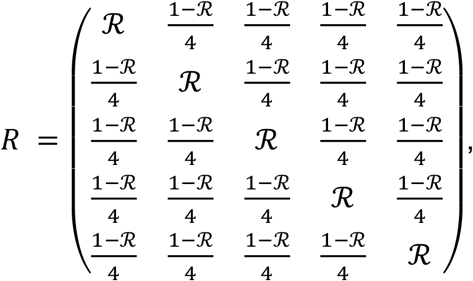

the death rates are 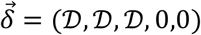, and the supply rates are, 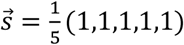 For 𝒟 =0.06 and ℛ = 0.85, the steady-state species abundances (from Eq. M.6) are 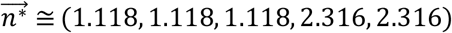) and steady-state resource concentrations (from Eq. M.5) are 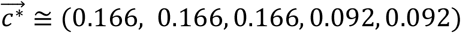 The corresponding two-species, two-resource community is represented by the consumption rate matrix

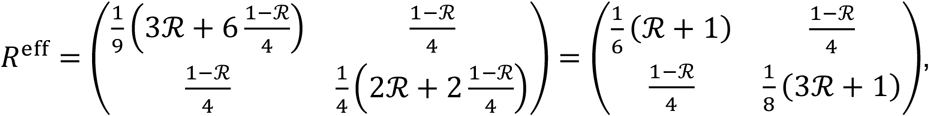

death rates 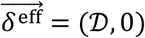 and supply rates 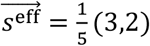 The steady-state species abundances and resource concentrations are 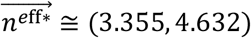 and 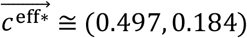 ,respectively. The sum of the steady-state abundances of species 1, 2, and 3 (with death rate 𝒟) equals the steady-state abundance of species 1 in the corresponding two-species, two-resource community. Similarly, the sum of the steady-state abundances of species 4 and 5 (with zero death) equals the steady-state abundance of species 2 in the corresponding two-species, two-resource community. A similar relation holds for the steady-state resource concentrations.

## Acknowledgements

We thank members of the Huang lab for helpful discussions. This work was funded by a Stanford Bioengineering Summer REU fellowship (to D.P.N.), a Stanford School of Medicine Dean’s Postdoctoral Fellowship (to P.H.), NIH Postdoctoral Fellowship F32 GM143859 (to P.H.), NSF Award EF-2125383 (to K.C.H.), and NIH Awards R01 AI147023 and RM1 GM135102 (to K.C.H.). K.C.H. is a Chan Zuckerberg Biohub Investigator.

## Code availability

Custom code used in this study can be downloaded at https://doi.org/10.5281/zenodo.13310240.

## Supplemental Figures

**Figure S1:**
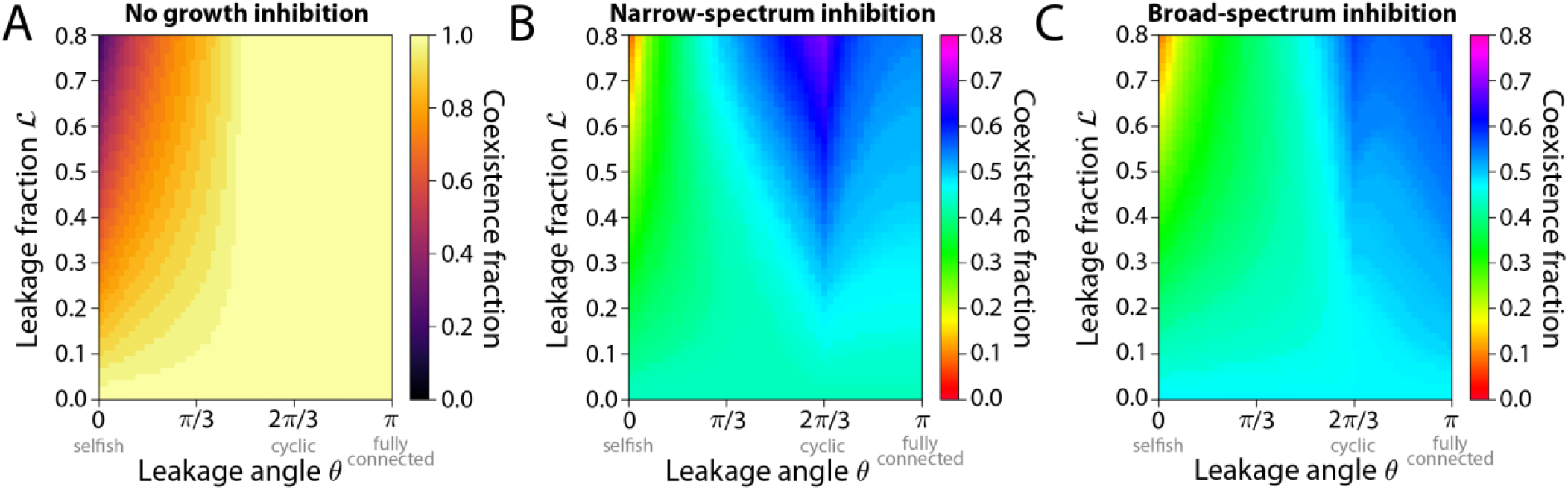
Inclusion of dilution has little effect on the conclusions of our study. Shown are re-creations of Fig. 2A (A), Fig. 3A (B), and Fig. 4A (C) without ignoring the dilution of resources term. Qualitative results were largely preserved.

**Figure S2:**
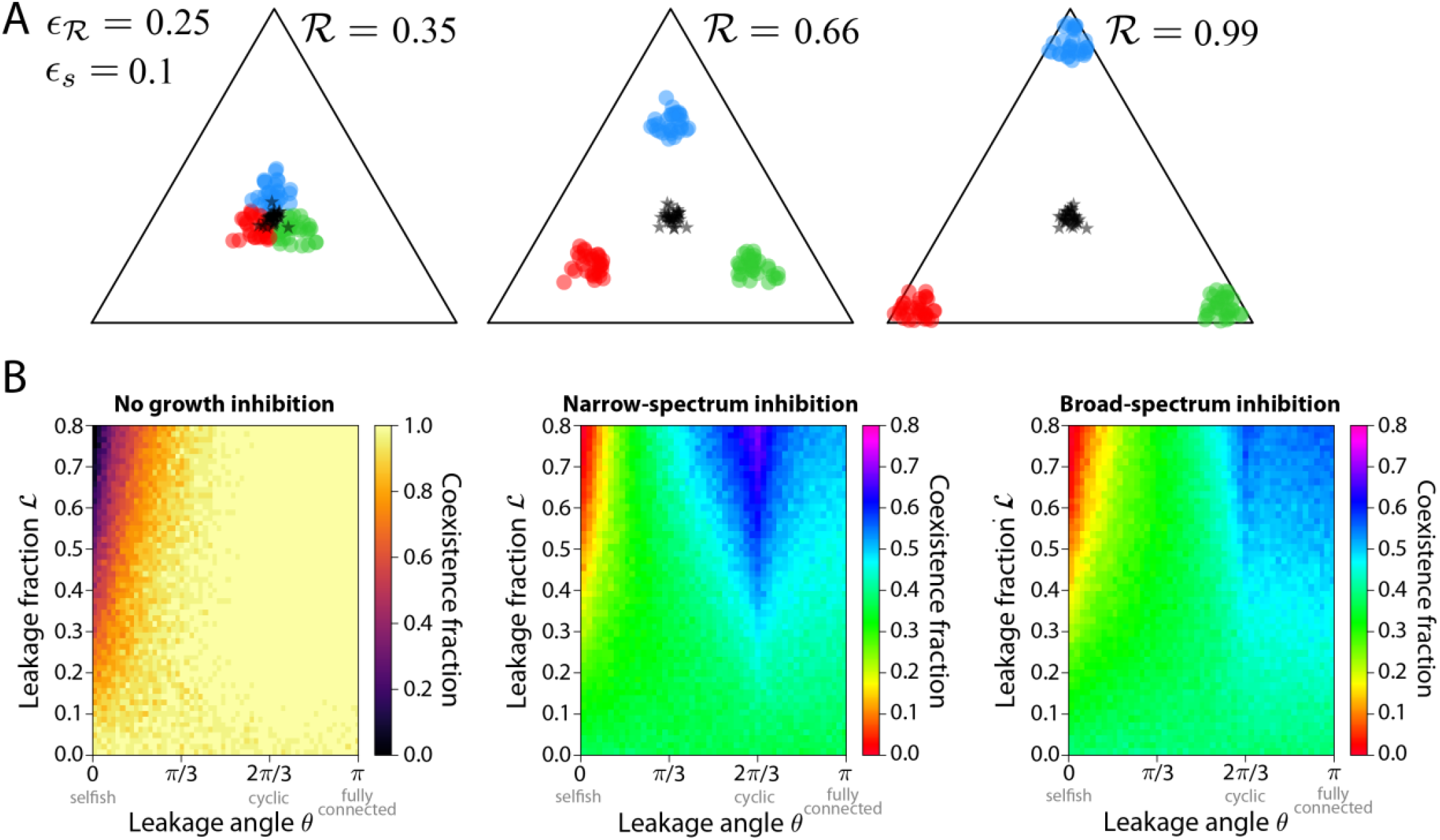
Our results were largely unchanged for non-uniform resource supply rates and non-symmetric resource consumption rates. A) We picked a vector 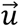 from a uniform distribution on the simplex and then replaced the resource supply rate vector 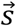 with a non-uniform resource supply rate that is the weighted average 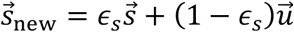, where *ϵ*_*s*_ is the extent of non-uniformity in resource supply rates. We similarly calculated new non symmetric resource consumption rate vectors in which the average was weighted by the extent of resource consumption symmetry breaking *ϵ*_ℛ_. For *ϵ*_*s*_ = 0.1 and *ϵ*_ℛ_ = 0.25, we visualized multiple instances of the randomly chosen resource consumption rates (black stars) and resource consumption rates (circles) for different resource consumption parameters ℛ. B) We repeated the calculations for the plots in Fig. 2A (left), Fig. 3A (middle), and Fig. 4A (right) but with randomly chosen supply rates and resource consumption rates as described in (A) and found that the results were largely unchanged.

**Figure S3:**
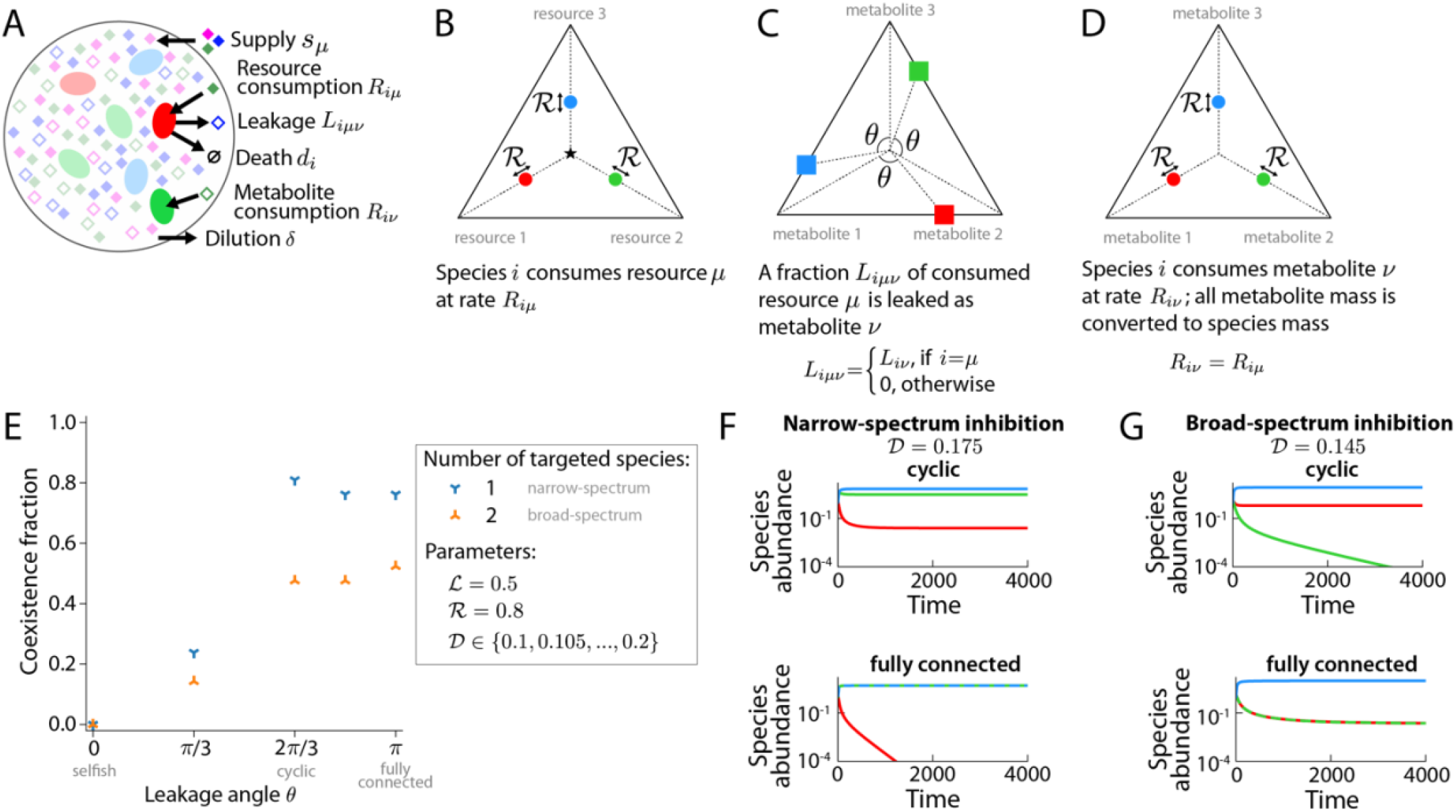
Modifying the CRL model to incorporate three primary nutrients supplied externally (resources) and three cross-fed nutrients not supplied externally (metabolites) does not affect the qualitative conclusions in Fig. 3A and Fig. 4A. A) Schematic of the model. Three externally supplied nutrients (solid diamonds) are consumed by three species (ovals) at rates given by the resource consumption rate matrix *R*_*iμ*_, which is the same as defined in the main text, in which each species prefers a unique resource. The fraction of resource *μ* consumed by species *i* that is excreted as metabolite *v* is *L*_*iμv*_; we assume that species *i* prefers to use *μ* = *i* for growth, and it uses resource *μ* = *i* to produce cross-fed nutrients, which effectively makes *L*_*iμv*_ a matrix (*L*_ii*v*_). The leakage fractions *L*_*iiv*_ is given by the matrix elements *L*_*iv*_ from the main text, and the metabolite consumption rates are given by the matrix elements *R*_*iμ*_, i.e., species *i* prefers to consume cross-fed metabolite *v* = *i*. B) Simplex depicting relative resource consumption rates (circles) as in Fig. 1B. C) Simplex depicting leakage angle *θ* and relative leakage rates of metabolites (squares). D) Simplex depicting relative metabolite consumption rates (circles). E) For total leakage fraction ℒ = 0.5 and extent of resource competition ℛ = 0.8, we calculated the coexistence fraction across death rates from 0.1 to 0.2 and found similar behavior as with the CRL model in the main text. Namely, for narrow-spectrum or broad-spectrum growth inhibition, the coexistence fraction is largest for cyclic or fully connected leakage, respectively. F) For death rate 𝒟 = 0.175 imposed on one species (mimicking a narrow-spectrum antibiotic), coexistence occurred with cyclic cross-feeding (top), but the targeted species (red) went extinct with fully connected cross-feeding (bottom). G) Like (F), but for death rate 𝒟 = 0.145 imposed on two species (mimicking a broad-spectrum antibiotic). One targeted species (green) went extinct with cyclic cross-feeding (top), but coexistence occurred with fully connected cross-feeding (bottom).

**Figure S4:**
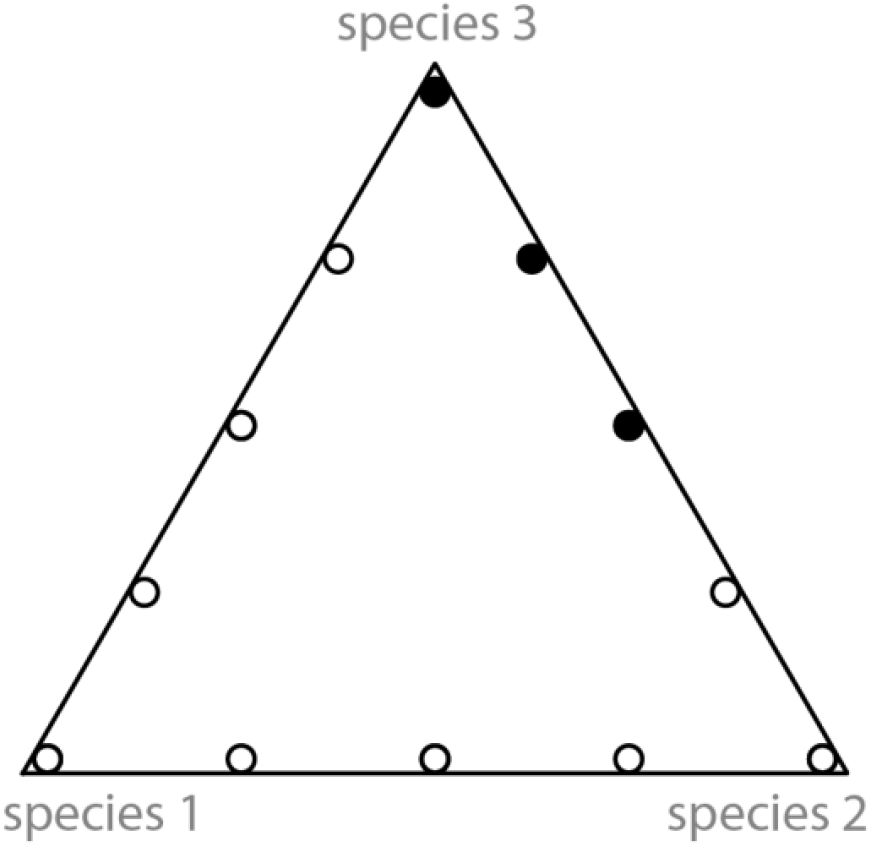
Simplex of relative initial species abundances used to test for multistability in randomly sampled communities in Fig. 2A. We calculated the population dynamics for three initial species relative abundances (filled circles). Because of the symmetry of communities in Fig. 2A, in which species labels are arbitrary, we effectively also tested nine other initial species relative abundances (hollow circles).

**Figure S5:**
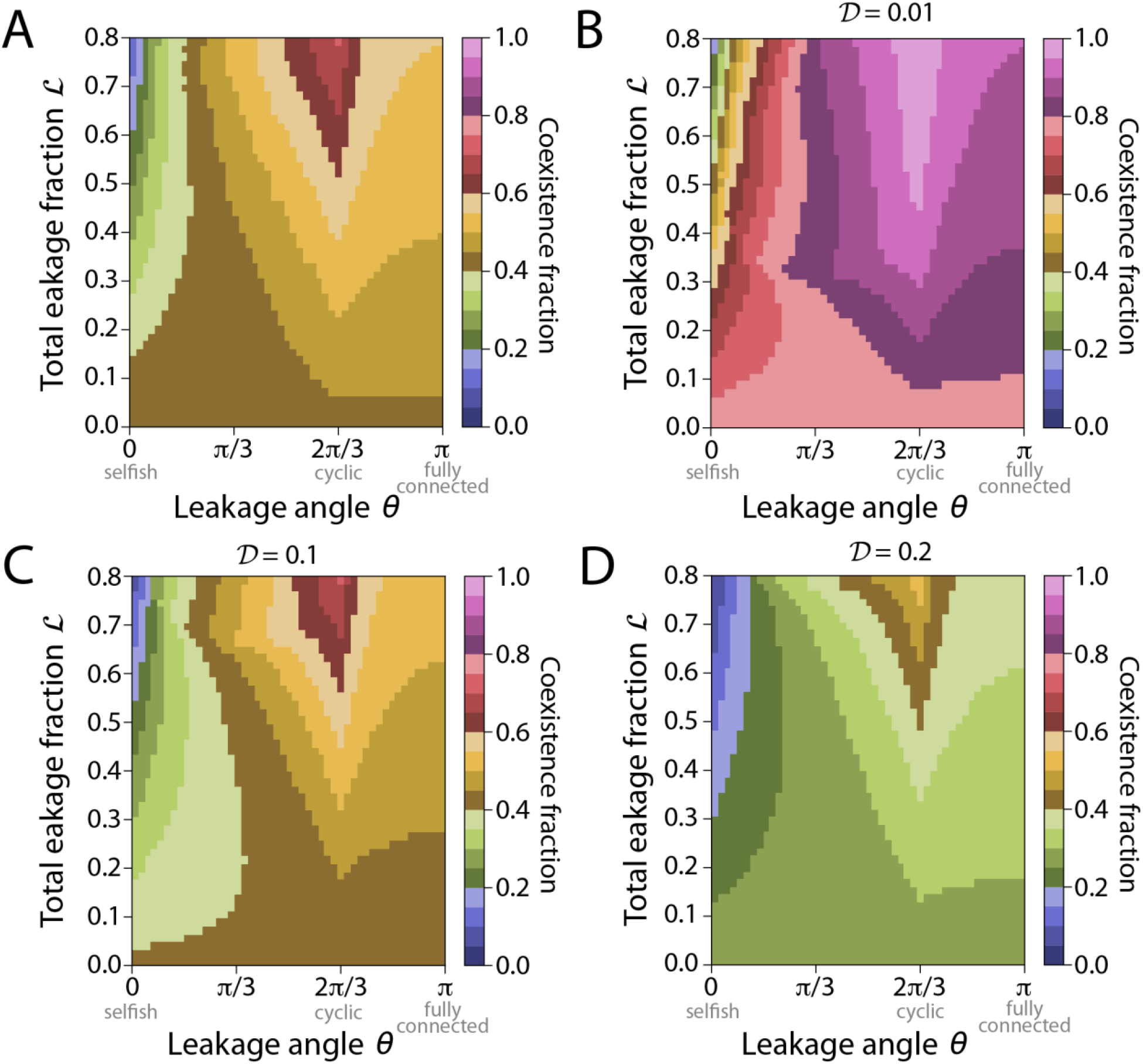
During narrow-spectrum stresses, coexistence is most likely with cyclic leakage regardless of death rate. A) The coexistence fraction across death rates 𝒟 ∈ [0.01, 0.2] and extent of resource competition ℛ ∈ [0.35, 0.99]; the plot is identical to Fig. 3A except with a different color bar for comparison with (B-D). B-D) The coexistence fraction averaged across extent of resource competition ℛ ∈ [0.35, 0.99] with fixed death rate 𝒟 = 0.01 (B), 𝒟 = 0.1 (C), or 𝒟 = 0.2 (D).

**Figure S6:**
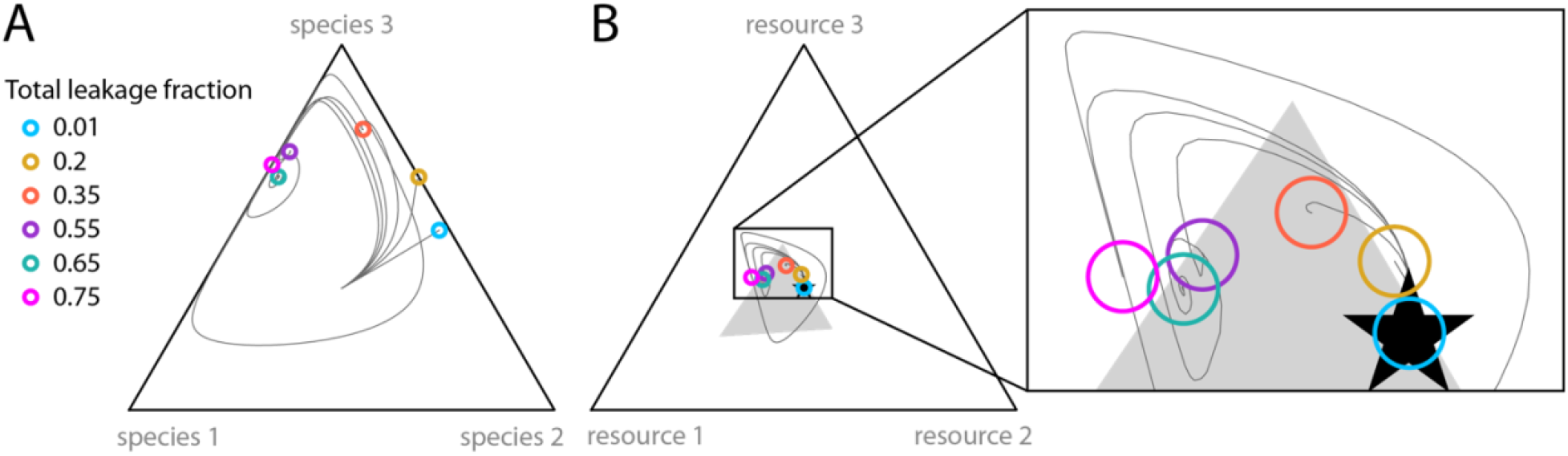
Richness changes non-monotonically with total leakage fraction during treatment with a narrow-spectrum stress. Population dynamics of relative species abundances (A) and relative effective resource supply rates (B) for simulations with six total leakage fractions (colored circles), resource competition ℛ = 0.5, leakage angle *θ* = *π*/4, and treatment with a narrow-spectrum stress with low death rate 𝒟 = 0.01 (*δ*_1_ = 0.01, *δ*_2_ = 0, *δ*_3_ = 0). Gray curves are the trajectories and colored circles represent the final time point of the simulation.

**Figure S7:**
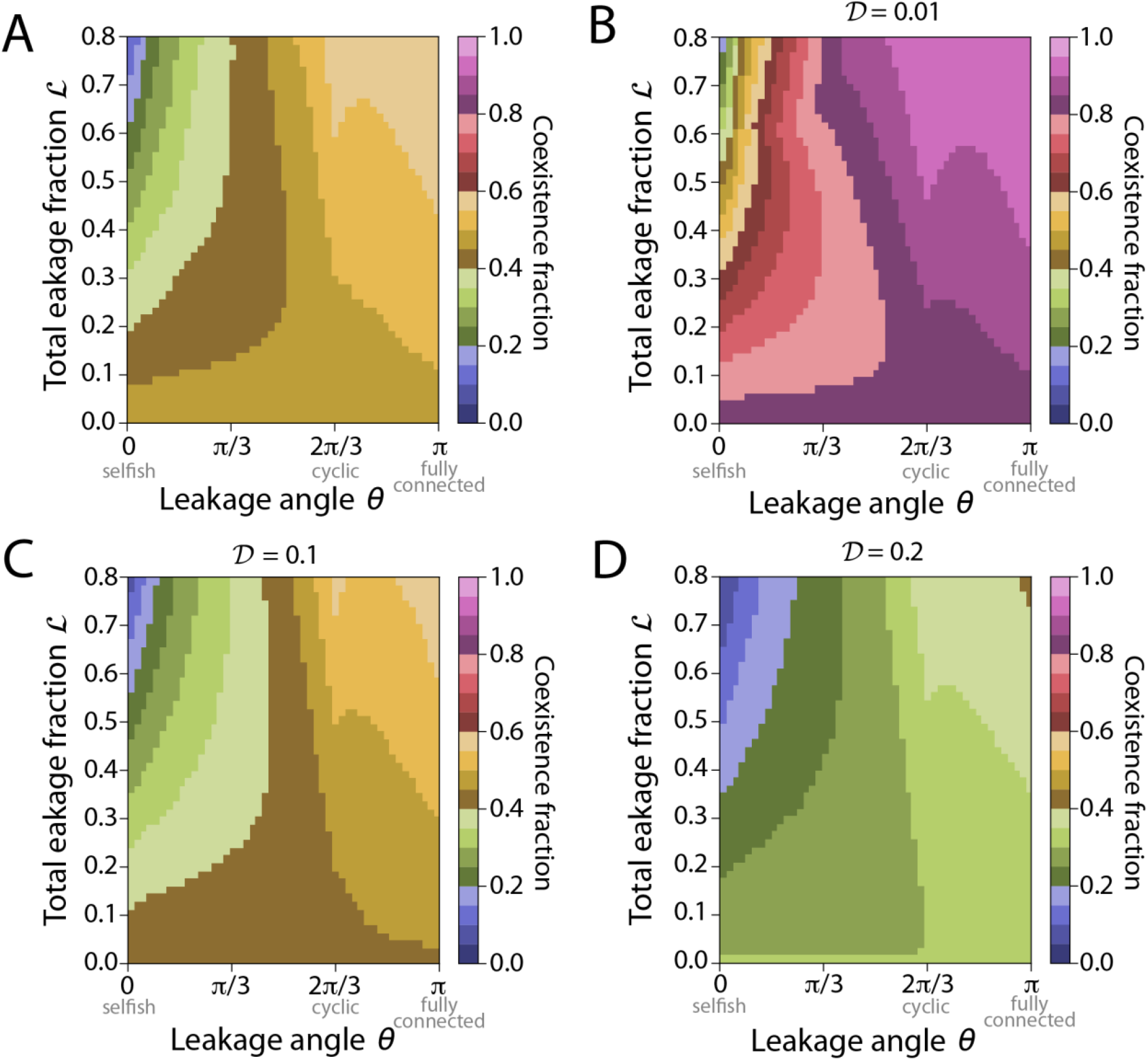
During a broad-spectrum stress, coexistence is most likely for fully connected leakage regardless of death rate. A) The coexistence fraction across death rates 𝒟 ∈ [0.01, 0.2] and extent of resource competition ℛ ∈ [0.35, 0.99]; this plot is identical to Fig. 4A except with a different color bar for comparison with (B-D). B-D) The coexistence fraction averaged across extent of resource competition ℛ ∈ [0.35, 0.99] with fixed death rate 𝒟 = 0.01 (B), 𝒟 = 0.1 (C), or 𝒟 = 0.2 (D).

**Figure S8:**
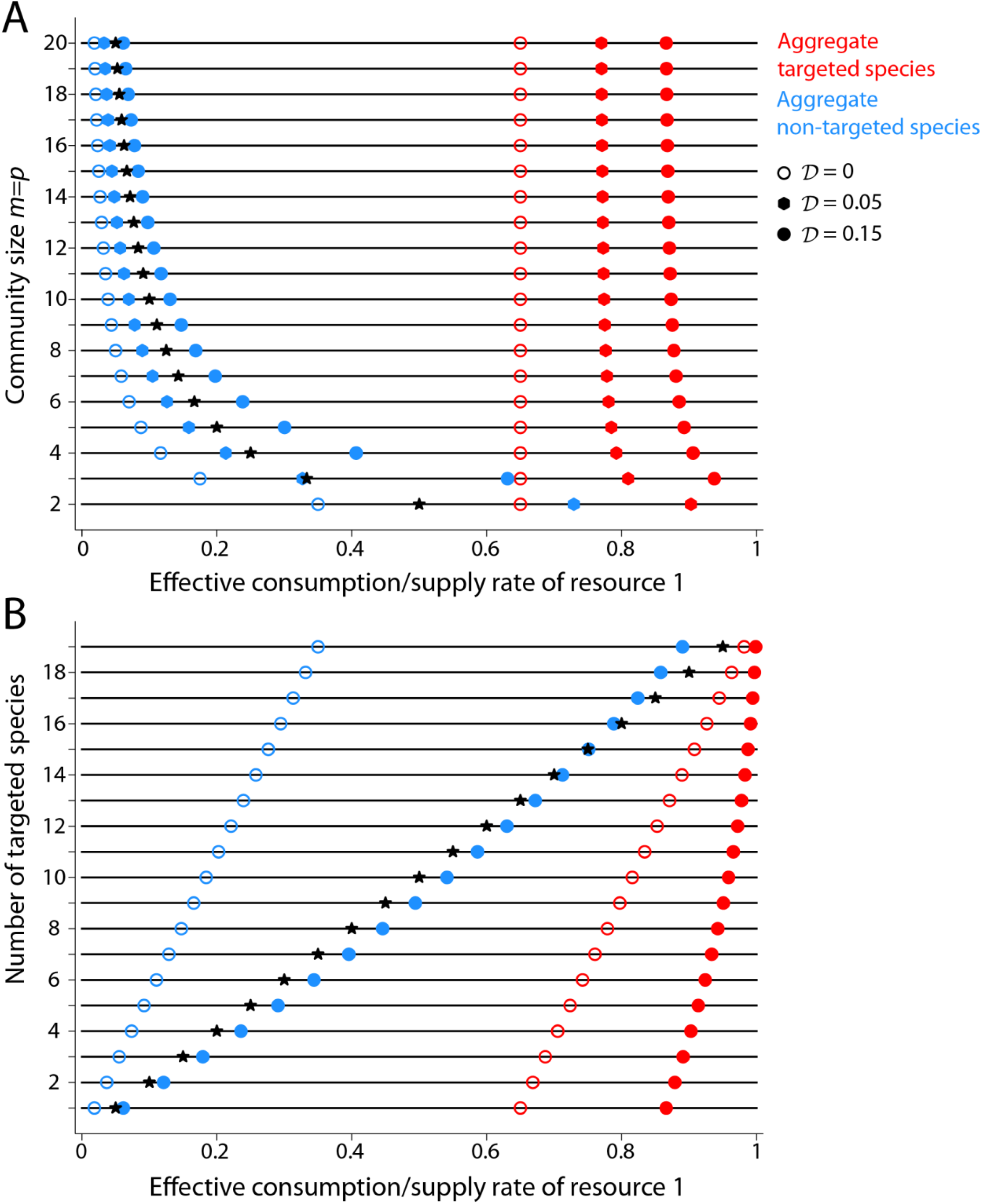
Coexistence region bounds and supply rates for an effective two-species, two-resource community shift as the actual community size or actual number of targeted species is increased. To visualize the coexistence region for larger community sizes, we exploited the symmetry among all non-targeted species given their resource consumption pattern to map them to a single effective non-targeted species and resource, respectively^22^ (Methods). A) As the actual community size increases, the relative supply rate (black stars) of the resource favored by the targeted species decreases since increasing the community size by 1 involves adding an additional equally supplied resource that is not favored by the targeted species. Without death (hollow circles), the consumption rate of the targeted species (red hollow circle) does not change as community size increases, but the consumption rate of the non-targeted species (blue hollow circle) decreases, indicating higher relative preference for its favored resource. When the death rate of the targeted species increases from zero, the coexistence region bounds shift toward the resource favored by the targeted species (filled octagons and circles). As community size increases, the supply rate decreases more slowly than the effective consumption rate of the non-targeted species (blue), hence the supply rate enters the coexistence region for larger death rates as the community size increases. B) Like (A), but for a community size *m* = *p* = 20 and a variable number of targeted species. When the number of targeted species is 1, the relative supply rate (black star) lie in the coexistence region when the death rate is zero (hollow circles), but not when 𝒟 = 0.15 (solid circles). When 𝒟 = 0.15, as the number of targeted species increases above 15, the supply rates enter the coexistence region (solid circles). Note that the case when the number of targeted species is 1 is the same as in (A) when the community size is 20.

